# Orchestration of differential mesodermal fate choice from ESCs by Wnt-USP3 link and H2A/H2B contextual deubiquitination

**DOI:** 10.1101/2023.07.01.547341

**Authors:** Varun Haran, Nibedita Lenka

## Abstract

Wnt, an evolutionarily conserved morphogen, is vital for various cell fates specification during early development. However, a concrete mechanistic understanding of the precise and fine-tuned regulation of Wnt underlying these processes is yet to be uncovered. Using the murine embryonic stem cells (ESCs) model, we have identified USP3, a histone deubiquitinase (DUB), displaying bimodal action, serving both as a downstream Wnt target and a regulator of canonical Wnt signalling. Using both loss- and gain-of-function approaches, we could identify USP3 as essential for mesoderm specification, exerting a differential influence during further differentiation. While cardiogenic mesoderm was negatively regulated and so also cardiomyogenesis, USP3 positively regulated hemangioblasts differentiation. Interestingly, however, these induced hemangioblasts promoted the haematopoietic program at the expense of endothelial differentiation. The mechanistic underpinning revealed USP3 localizing to chromatin and differentially modulating these fate choices by precise and contextual deposition of H2AUb/H2BUb in the promoters of mesoderm genes. Collectively our study underscored the Wnt-USP3 link underlying differential mesodermal fate modulation.

## Introduction

Vertebrate embryogenesis is highly complex, involving well-orchestrated developmental events that are tightly regulated by a cascade of various signalling pathways and their cross-talk. Indeed, both genetic and epigenetic modulators exert their temporo-spatial influence governing the intricacies during development. Embryonic stem cells (ESCs) that serve as an elegant *in vitro* developmental model effectively recapitulate early embryogenesis during differentiation and possess the potential to generate multiple mesodermal derivatives in culture in response to specific morphogens^1^. Several reports including ours entail the crucial influence of Wnt, activin/Nodal, Cripto, bone morphogenetic proteins (BMPs), fibroblast growth factors (FGFs), Hedgehog, Notch, etc. during mesoderm induction and cardiogenesis^2–6^. Wnt/β-catenin signalling induces the specification of hemangioblast population that later drives both haematopoietic and endothelial cells’ differentiation^7–9^. However, Wnt response with respect to cardiac development and cardiomyocyte differentiation remains contentious due to several contradictory reports in different model organisms, and also based on the conditions provided during *in vitro* differentiation^10–13^. Even, a temporal influence of Wnt during mesodermal induction from ESCs has been reported by us demonstrating canonical Wnt activation during the early stage of differentiation helping in the induction of mesoderm and endoderm, in contrast to attenuating cardiomyogenesis^4^. This brings up an interesting paradigm concerning stage-specific Wnt signalling modulation by various genetic and epigenetic cues.

Protein ubiquitination, an event that regulates the signalling threshold functioning as a signalling intermediate, targets the proteins for degradation and it has also been implicated in Wnt signalling modulation^14^. The signalling events in the Wnt cascade intersect with the turnover of its transcriptional activator, β-catenin, by ubiquitin-proteasome mediated degradation^14^. Moreover, there exists deubiquitination, conferred by deubiquitinating enzymes (DUBs), to counterbalance the ubiquitination process in a reversible manner^15^. Hence, maintenance of this intricate balance is essential in dictating the signalling threshold, expression of enzymes and structural proteins, and thereby guiding the cell fate decision machinery. Several independent studies reveal the involvement of ubiquitin-specific proteases (USPs), like USP4, USP6, USP7, USP14, USP22, and USP47 in regulating canonical Wnt signalling^16–21^. Similarly, USP12, USP16, USP22, and USP44 have been reported to be important for mesoderm development and interestingly, all these are associated with deubiquitinating histones H2A and/or H2B^22–25^. However, none of these studies have demonstrated DUBs that are involved during mesoderm differentiation as the effectors of canonical Wnt signalling or *vice versa*. A rigorous computational search for these DUBs has led us to the identification of USP3, a reported histone deubiquitinase, as a Wnt target gene during mesoderm differentiation.

Previous studies had documented USP3 deletion leading to increased histone ubiquitination in adult tissues and its importance in maintaining HSCs homeostasis by preserving HSCs self-renewal and repopulation potential and also in DNA damage mechanisms^26–28^. Here, using the murine ESCs model, we have demonstrated a novel role of USP3 displaying a bimodal action, both as a target and a regulator of canonical Wnt signalling. Our data have revealed USP3 mediates the fate-switching between neuroectoderm and mesoderm specification from ESCs by attenuating the former and promoting the latter. Moreover, USP3 exerts temporal and pronounced effects during further differentiation into various mesodermal derivatives. The mechanistic underpinning has elucidated USP3 deubiquitinating both H2AUb and H2BUb and their differential deposition underlying the regulation of various mesodermal genes expression. Overall, our study has suggested a context-dependent, both positive and negative modulatory influence of USP3, a Wnt target, during mesodermal fate choice and to its derivatives.

## Results

### USP3 is a conserved Wnt target during Mesoderm Differentiation

Wnt is well proven as an indispensable player for mesoderm specification during embryonic development^4, 29–32^. In concurrence to that, quantifying Brachyury (Bry) expressing cells by flow cytometry during mesodermal differentiation from ESCs revealed Wnt activation inducing mesoderm specification, while its inhibition suppressing the same (Fig. 1a). To delineate the mechanistic basis underlying Wnt influence on mesoderm differentiation, *in silico* analysis was carried out on retrieved RNA sequencing data deposited in Gene expression Omnibus (GEO) platform^33, 34^ for identifying plausible players. Our search using “GREIN”, a GEO RNA-seq data analyzing platform^35^ narrowed it down to USP3, a DUB, as a crucial candidate that was seen up-regulated during mesoderm differentiation. The mouse gene expression database during embryonic development (GXD) also revealed high expression of USP3 among mesodermal derivatives^36^. Interestingly, the promoter elements of USP3 contained conserved TCF/LEF binding motifs, hence prompting us to verify the USP3-Wnt link during mesoderm differentiation. ChIP analysis revealed β-catenin recruitment to the promoter region of USP3 upon active Wnt signalling [∼ 8-fold with BIO^4^ (6-bromoindirubin-3′-oxime), a GSK-3β inhibitor, when compared to DMSO] during mesoderm differentiation (Fig. 1b, b’), and thus illustrated USP3 as a novel Wnt target gene. Likewise, Wnt activation also induced *USP3* gene expression by 3-5 folds, when analyzed on d2, d5, and d10 of ESCs differentiation to cardiomyogenic lineage (Fig. 1c). This signified USP3 as a downstream effector molecule of canonical Wnt signalling during mesoderm specification and further differentiation. Moreover, the peak expression of *USP3* was seen on d5 of ESCs differentiation, the stage coinciding with that of optimum *Bry* expression reported by us earlier^4^. Together these results underscored a plausible role of USP3 as a target of β-catenin-dependent canonical Wnt signalling during mesoderm specification and subsequent differentiation to its derivatives.

**Fig. 1:**
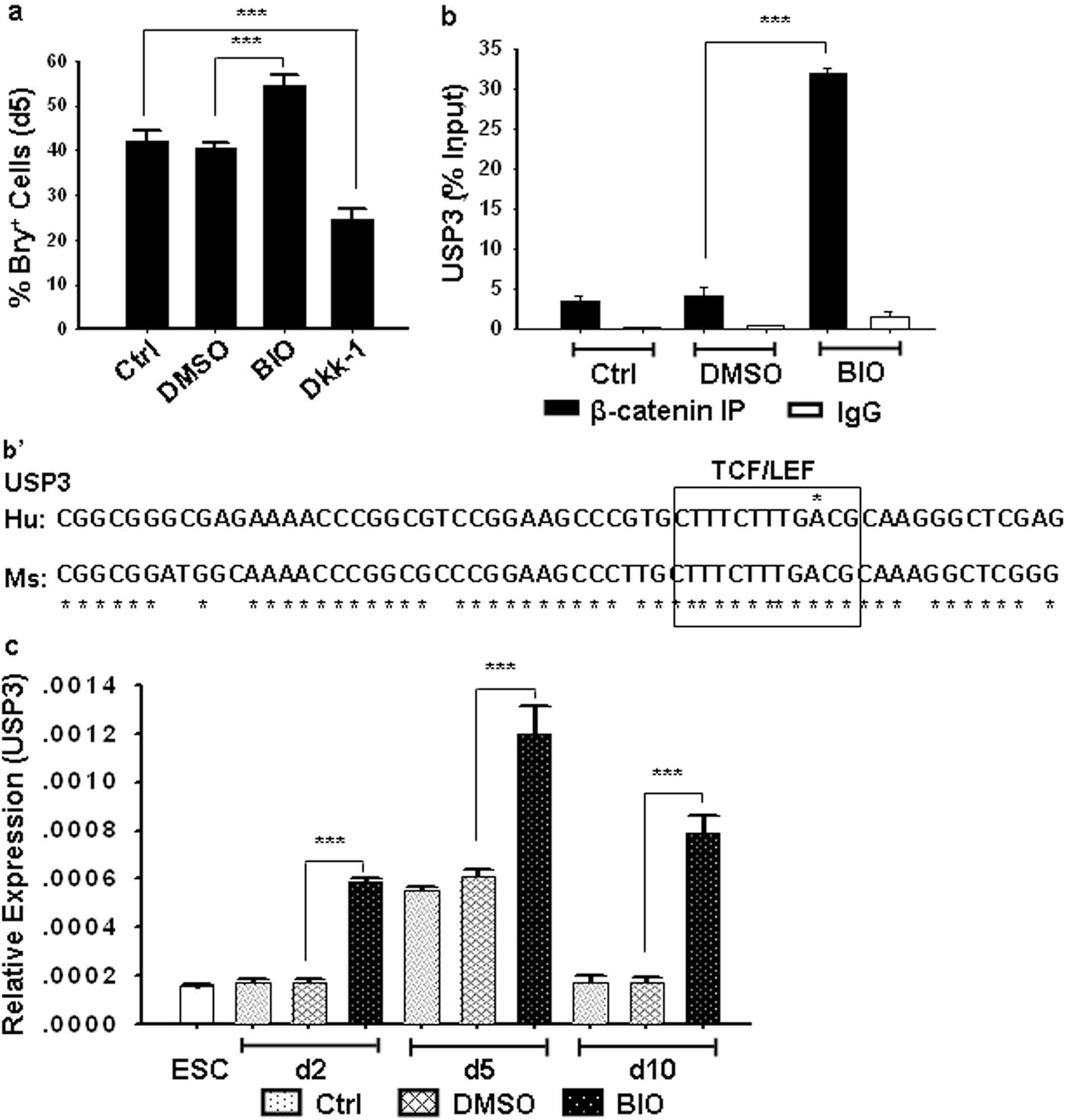
USP3, as a novel Wnt target, drives Mesoderm Differentiation from ESCs in vitro. (a) Flow cytometry quantification of cells on d5 of ESCs differentiation revealed an increased population of Bry^+^ cells upon BIO treatment compared to control, while Wnt inhibition with Dkk-1 attenuated the same; n = 4. (b) Wnt activation could increase the occupancy of β-catenin to the USP3 promoter region (-180 to +60 bp) as discerned by chromatin immunoprecipitation followed by qPCR. While IgG was used as a negative control, sonicated genomic DNA served as the input control; n = 3. (b’) USP3 promoter is having conserved sequences resembling TCF/LEF binding motif closer to the transcription start site (human: -9/+3; mouse: -46/-35); sequence retrieved from Eukaryotic Promoter Database and alignment carried out using CLUSTAL ω (1.2.4). (c) qPCR showing an increase in USP3 expression with Wnt activation on d2, d5, and d10 of ESCs differentiation compared to undifferentiated ESCs and the respective DMSO controls. Further, maximum expression of USP3 was observed on d5 of mesoderm differentiation. Data are means ± SEM; n = 3-4. *, P ≤ 0.05; **, P ≤ 0.01; ***, P ≤ 0.001.

### USP3 does not alter ESC characteristics during maintenance

Next, we evaluated whether USP3 would also regulate ESC characteristics during maintenance similar to Wnt^37^. We generated both USP3 deficient (USP3-KD) and efficient (USP3-O/E) stable ESC clones along with its catalytic mutant (USP3-mt) clones to assess the functional attribute of USP3 and its catalytic domain in rendering Wnt responsive action. The validation for the stated clones was carried out at both transcriptional and translational levels (Fig. 2a, b). Neither USP3-KD nor USP3-O/E clones exhibited any appreciable difference with respect to either the colony morphology (Fig. 2c) or the SSEA1^+^ population (∼90%) when compared with their respective control counterparts. The same was also true with the expression of other key undifferentiated ESC markers (Oct4, Nanog, Sox2, cMyc, Klf4) in them (Fig. 2d-f, Fig. S1a-d). Similarly, withdrawing LIF and maintaining these cells for 5 days in culture resulted in spontaneous differentiation with reduced expression of pluripotency associate marker genes similar to that seen in control (Fig. S2). Together, it was inferred that USP3 might be dispensable during ESCs’ self-renewal and retention of pluripotent characteristics in them.

**Fig. 2.**
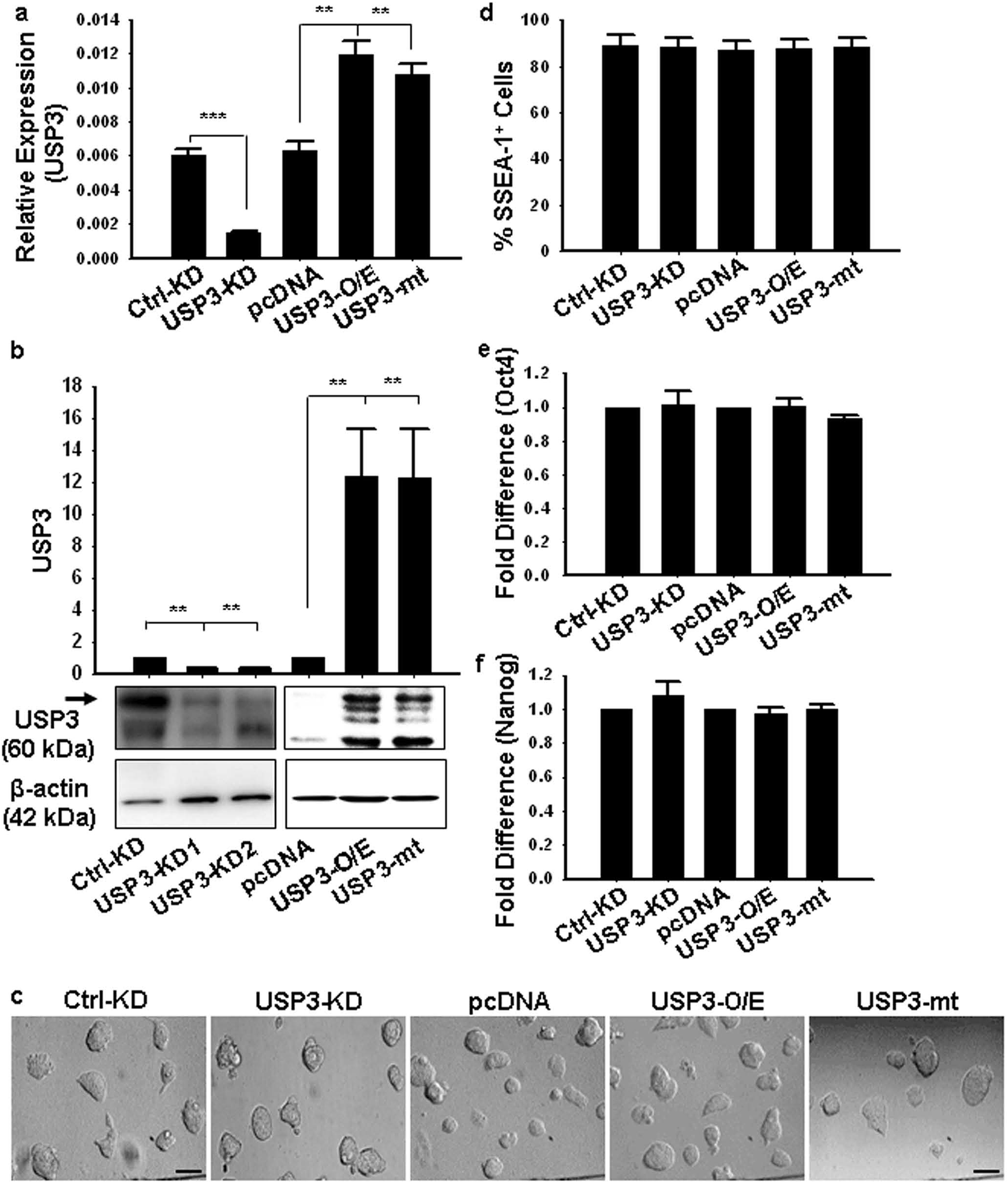
USP3 is dispensable during ESC maintenance. (a) qPCR analysis showing a reduction in *USP3* expression in USP3-KD ESCs, while the same was increased in USP3-O/E and USP3-mt clones compared to their respective controls; n = 3. (b) Western blot analysis showing a decrease in USP3 protein levels with USP3-KD and the same being increased with USP3-O/E and USP3-mt clones compared to their respective controls. Data were presented as fold differences keeping the respective control value as 1; n = 4-6. (c) Bright-field images of USP3-KD, USP3-O/E, and USP3-mt cells showing similar colony morphology under the maintenance condition as that of respective control cells. (d) Flow cytometry quantification of SSEA1^+^ ESCs during maintenance revealed no significant difference among the stated clones; n = 4. (e,f) Quantification of *Oct4* (e) and *Nanog* (f) transcripts by qPCR during ESCs maintenance showed no significant difference among the clones irrespective of USP3 status. Data were presented as fold differences keeping the respective control value as 1; n = 3. Data are means ± SEM; **, *P* ≤ 0.01; ***, *P* ≤ 0.001.

### USP3 promotes Mesoderm induction while blocking neurectoderm differentiation from ESCs

To further assess USP3 induced-differentiation status in ESCs, whether uniformly to all three lineages or preferentially to any specific one during spontaneous differentiation, expression of three germ layer-specific markers, such as Nestin (Nes) (ectoderm), Bry (mesoderm), and α-feto protein (AFP) (endoderm) was analyzed by immunocytochemistry in cells cultured for five days under -LIF conditions. USP3-KD specifically promoted Nes expression, inhibited Bry, and had no influence on AFP (Fig. 3a). The reverse was true in the case of USP3-O/E. The same was further verified at the transcriptional level that showed ∼ 3-fold induction and 2-fold reduction in *Nes* and *Bry* expression respectively in USP3-KD compared to control (Fig. 3b), and hence, suggested its differential influence on germ layer specification from ESCs *in vitro*. Monitoring the *in vivo* pluripotency through teratoma assay also revealed strikingly higher Nes^+^ cells as well as Nes expression (intensity) in contrast to reduced Bry expression and overall Bry^+^ cells in the USP3-KD group compared to control and the reverse being true in USP3-O/E group (Fig. S3). Thus, USP3-induced differentiation in ESCs displayed an interesting differential lineage fate-switching event between the mesoderm and the neuroectoderm, and the endoderm remained unaffected.

**Fig. 3:**
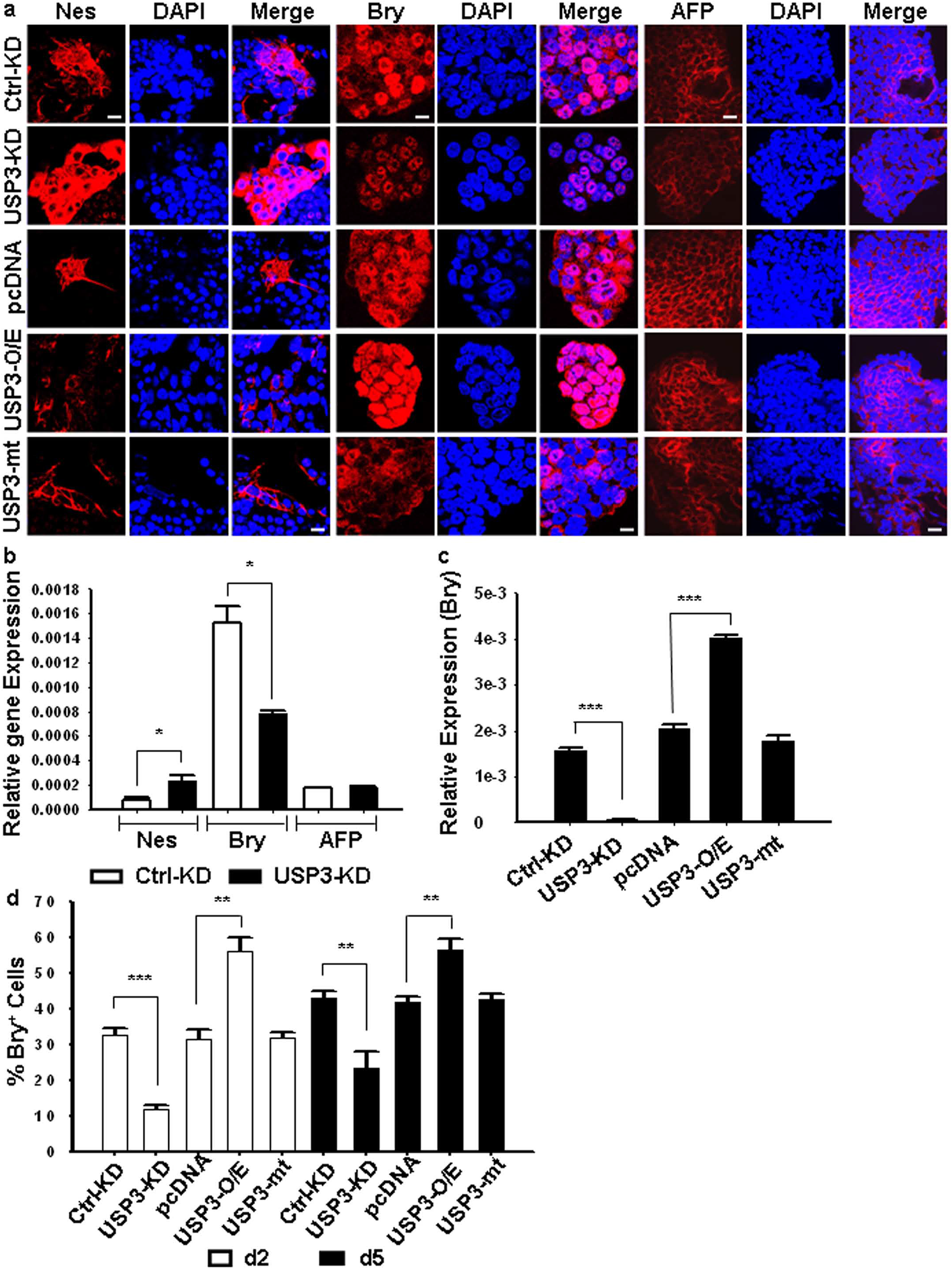
USP3 exerts differential influence during induction of mesoderm and neurectoderm from ESCs *in vitro*. (a) Immunofluorescence analysis for Nestin (Nes), Brachyury (Bry), and AFP during spontaneous differentiation of stated ESC clones maintained under -LIF condition for 5 days. USP3 imparted a negative regulatory influence on neuroectoderm induction in contrast to positive regulation with respect to mesoderm induction. Scale bar: 20μm. (b) Expression status of *Ne*s, *Bry,* and *AFP* transcripts during spontaneous differentiation of stated ESC clones maintained under -LIF condition for 5 days. (c) Expression status of *Bry* on d5 of mesodermal differentiation from ESCs *in vitro* revealed its decreased expression in USP3-KD, while the same was increased in USP3-O/E clone when compared with their respective controls; n= 3. (d) Flow cytometry quantification of Bry^+^ cells on d2 and d5 of ESCs differentiation validated a mesoderm promoting effect of USP3; n = 3-4. Data are means ± SEM; *, P ≤ 0.05; **, P ≤ 0.01; ***, P ≤ 0.001.

Next, we proceeded with directed differentiation to gain further insight into the influence of USP3 on cell fate modulation especially to mesodermal lineage and its derivatives. While USP3-KD led to decreased expression in *Bry* (∼ 27-fold) compared to control, the same was seen up-regulated in USP3-O/E (∼ 2-fold), but not in USP3-mt, when monitored on d5 of ESCs differentiation (Fig. 3c). Flow cytometry quantification also revealed USP3-KD decreasing the Bry^+^ cells when monitored on both d2 and d5 of ESCs differentiation compared to that in control (Fig. 3d). In contrast, the same was seen increased in USP3-O/E as early as d2 of differentiation. However, a similar % of Bry^+^ cells obtained in the case of both USP3-mt and control did indeed suggest the involvement of USP3 catalytic activity in positively modulating mesoderm specification and expediting the same during ESC’s differentiation.

Previous studies have shown that the *Bry* gene is a direct target of Wnt signalling, which recruits the Wnt transcriptional effectors such as β-catenin/TCF/LEF, important for mesoderm patterning, to its promoter^29, 38^. Considering *Bry* expression is at its peak on d5 of ESCs’ differentiation^4^, we assessed whether USP3 might as well modulate Wnt signalling during mesoderm differentiation, with the readout being monitoring the levels of β-catenin at the specified time regimen. Immunoblotting analysis on d5 EBs using USP3 deficient and efficient clones revealed USP3-KD decreasing the expression of β-catenin (∼ 2-fold), while USP-O/E increasing it (Fig. 4a). Similar trend was also seen at the transcriptional level (Fig. 4b) suggesting USP3 might be possibly modulating Wnt signalling by either regulating the expression of β-catenin or preventing its proteasomal degradation. Further, validation was obtained upon detection of nuclear β-catenin, reflecting active Wnt signalling under USP3 efficient condition (Fig. 4c). Indeed, the direct interaction between β-catenin and USP3 through co-immunoprecipitation studies (Fig. 4d, e) unequivocally strengthened the canonical Wnt-DUB link underlying mesodermal specification and differentiation. Moreover, this together with the data showing USP3 as a Wnt target (Fig. 1) brought an interesting paradigm of reciprocal regulation of Wnt and USP3 by each other and hence evoked interest in delineating the precise role of USP3 during further differentiation into mesodermal derivatives.

**Fig. 4.**
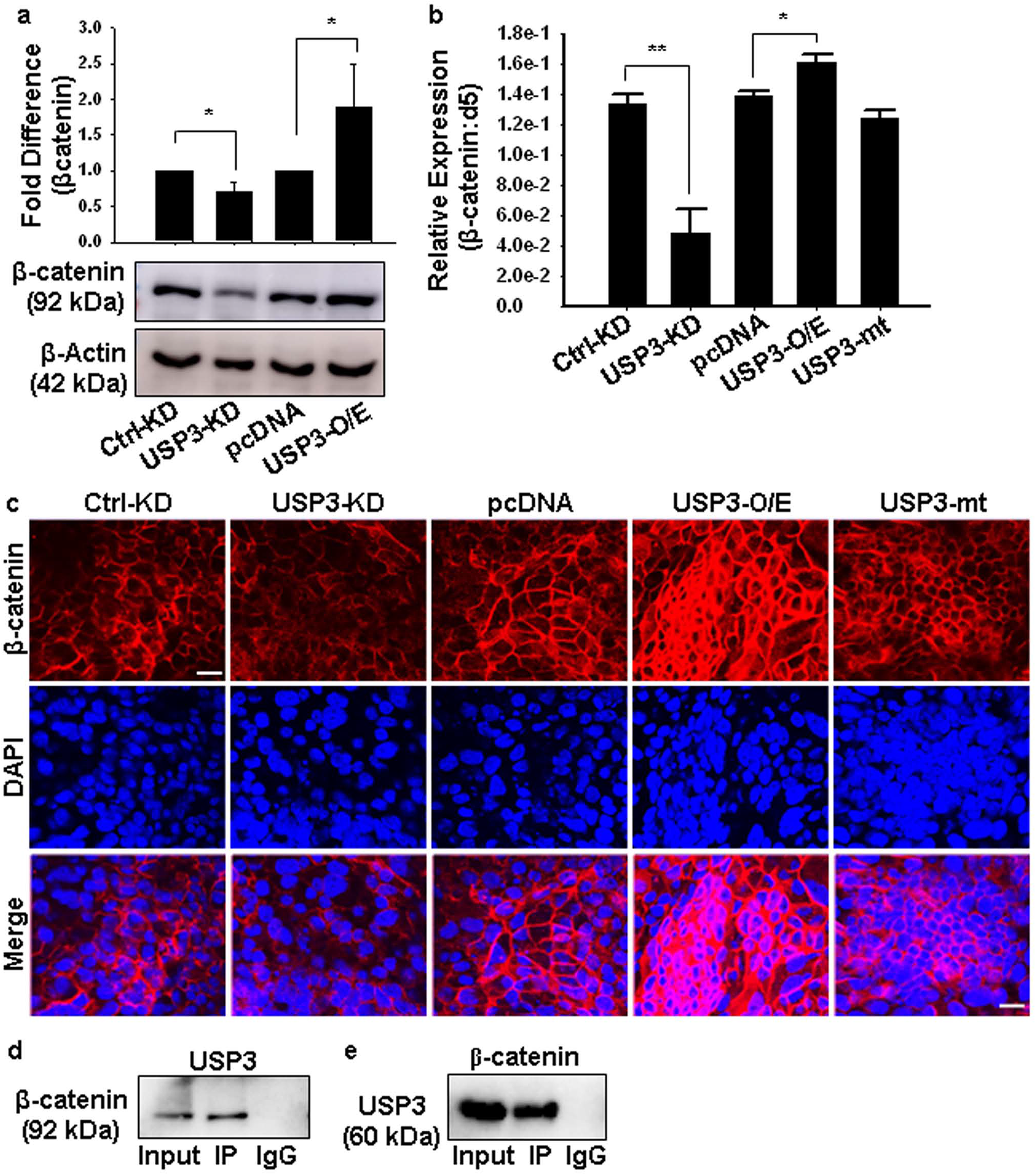
Reciprocal regulation of Wnt and USP3 during mesoderm differentiation (d5 EBs). (a) Western blot analysis revealed β-catenin expression decreasing upon USP3 knock-down (USP3-KD), while the reverse was true in case of its overexpression (USP3-O/E) when compared with their respective controls; n = 4. (b) qPCR analysis illustrated a positive influence of USP3 on *β-catenin* expression; n = 3. (c) Immunofluorescence analysis for β-catenin displaying its decreased expression in USP3-KD clone with respect to control, while the same was enhanced in USP3-O/E when compared to pcDNA or USP3-mt expressing clones. An increase in the immunopositivity of β-catenin in the nucleus upon USP3-O/E indicated the activation of canonical Wnt/β-catenin signalling. DAPI (blue) was used for nuclear staining, Scale bar: 20μm. (d.e) Immunoprecipitation using d5 EB cell lysates demonstrated USP3 interaction with β-catenin. The protein samples were immunoprecipitated with anti-USP3 or anti-β-catenin, followed by immunoblotting with anti-β-catenin or anti-USP3, respectively; n = 3. Data are means ± SEM; *, P ≤ 0.05; **, P ≤ 0.01.

### USP3 negatively regulates Cardiac progenitor specification and Cardiomyogenesis

Bry^+^ mesoderm attains two alternate cell fates, *i.e.*, Mesp1^+^ cardiac Mesoderm and Er71^+^ hemangioblasts during mesoderm differentiation^39–42^. Hence, we monitored the role of USP3 during cardiac mesoderm specification under both USP3 deficient and efficient conditions using the Mesp1-Venus ESC clone. While >50% of the cells were Mesp1-Venus^+^, ∼ 60% of the cells were PDGFRα^+^ in USP3-KD, and the same decreased to ∼ 30% and 27% respectively in USP3-O/E when monitored on d5 of ESCs’ differentiation (Fig. 5a-c, Fig. S4a). Interestingly, a similar trend was also observed as early as d2 of ESCs’ differentiation (Fig. 5a, USP3-mt clones however did not alter the population of either Mesp1 or PDGFRα compared to control on both days, reflecting thereby the deterministic influence of USP3 activity on cardiac mesoderm specification. These findings were further replicated by analyzing *Mesp1* expression on d5 of ESCs’ differentiation that revealed ∼ 2-fold increase and ∼ 7-fold decrease in USP3-KD and USP3-O/E respectively when compared with their respective controls and USP3-mt (Fig. 5d). Taken together, our data revealed USP3 having a negative modulatory influence on the generation of multipotent cardiac progenitors.

**Fig. 5.**
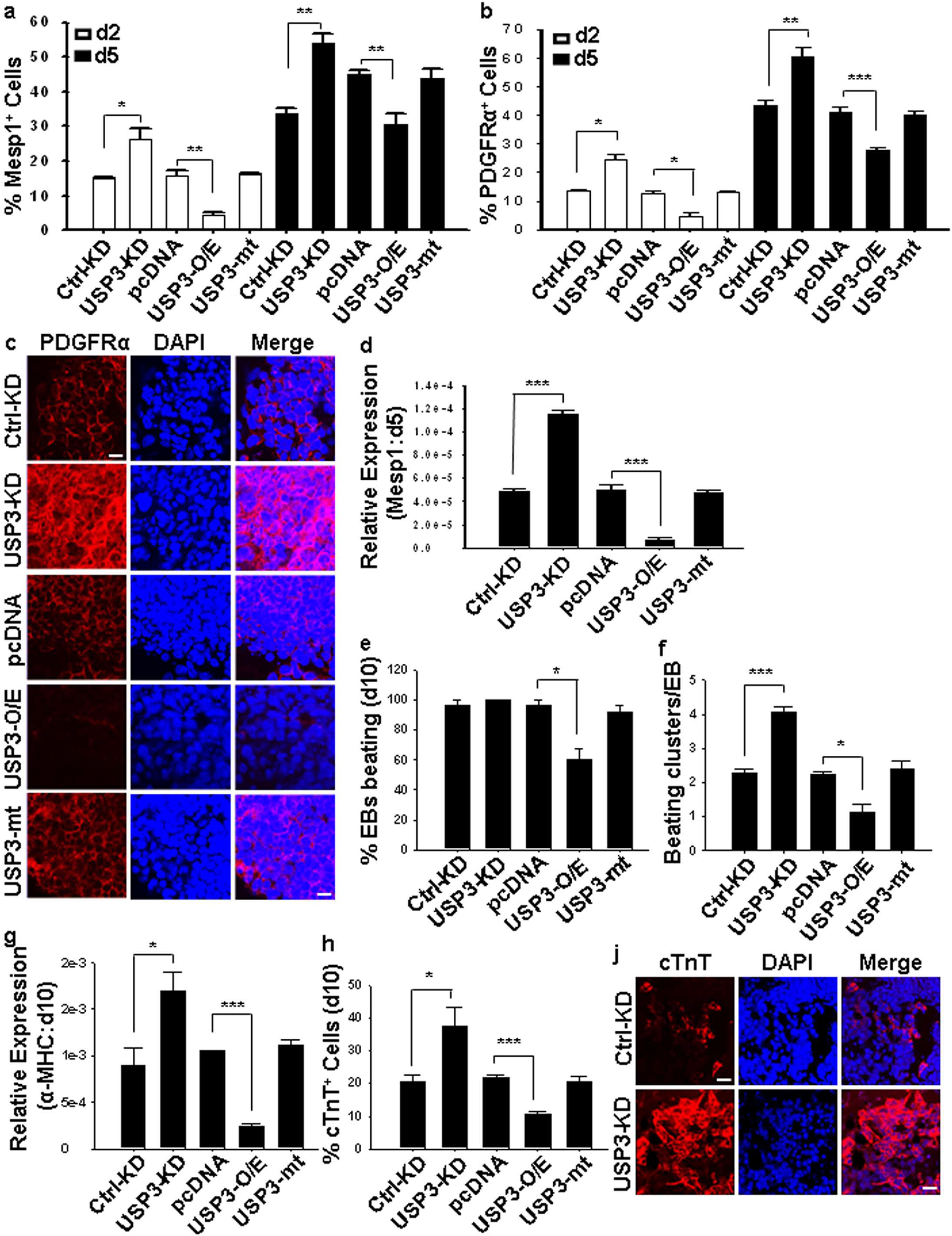
USP3 negatively regulated cardiac progenitor generation and cardiomyogenic differentiation. (a,b) Flow cytometry quantification of (a) Mesp^+^ and (b) PDGFRα^+^ cells in USP3 deficient and efficient clones using d2 and d5 EBs revealed a negative modulatory effect of USP3 on cardiac mesoderm specification. (c) Immunofluorescence analysis for PDGFRα^+^ cells on d5 of ESC differentiation also revealed a negative influence of USP3 on the generation of cardiac progenitors. Scale bar: 20μm. (d) qPCR analysis for the expression of *Mesp1* in d5 EBs revealed an increase in its expression with USP3-KD while decreasing the same with -O/E when compared with their respective controls. (e,f) USP3 attenuated cardiomyogenesis as quantified by counting the percentage EBs beating (e) and beating clusters/EB (f) on d10 of ESCs differentiation; n = 3-7). (g) qPCR analysis for α-MHC in d10 EBs showed an increase in its expression with USP3-KD while decreasing the same with -O/E and no change with USP3-mt when compared with their respective controls; n = 3). (h) Flow cytometry quantification of cTnT^+^ cells on d10 of cardiomyogenic differentiation of ESCs revealed a negative modulatory effect of USP3 on cardiomyogenesis; n = 3. (i) Immunofluorescence analysis for cTnT^+^ with USP3-KD during d10 of ESCs differentiation showed an increase in its generation. Quantitative data are means ± SEM; *, P ≤ 0.05; **, P ≤ 0.01; ***, P ≤ 0.001.

Consequently, to discern the role of USP3, the Wnt target, in cardiac differentiation regulation, USP3-KD and Ctrl-KD ESCs were differentiated and cardiac differentiation efficiency was compared by quantifying the % EBs beating, and counting the number of beating clusters/EB on d10 of ESCs’ differentiation (Fig. 5e, f). USP3-KD clones displayed enhanced cardiac differentiation potential by exhibiting increased beating clusters/EB compared to control and expressing a higher level of cardiac-specific genes; *α-MHC*, *Mef2c*, *Mlc2a* (Fig. 5f, g; Fig. S4b, c). Flow cytometry quantification and immunostaining also revealed USP3 deficiency increasing the generation of cardiac troponin (cTnT^+^) cells as well as its expression, compared to that of control (Fig. 5h, j). In contrast, USP3-O/E showed a reduction in cardiac differentiation efficiency as discerned by the afore-stated criteria (Fig. 5e-h). Collectively, our data suggested USP3 negatively regulated cardiac differentiation from mesoderm during ESC’s differentiation.

### USP3 positively regulates Hemangioblasts specification and differentially influences haemato-endothelial differentiation

Flk1^+^ mesoderm/hemangioblasts are derived from Bry expressing mesoderm, and Er71 that functions as a downstream Wnt target regulates the generation of these and also their derivatives such as haematopoietic and endothelial cells^39, 40, 42^. Considering USP3 dictated a contrasting modulation on mesoderm induction (positive) and subsequent cardiomyogenic differentiation (negative) programme, it became imperative to investigate whether or not the induced mesoderm could opt for an alternate cell fate such as hemangioblast in response to USP3 activation. Interestingly, USP3 displayed a temporal response on hemangioblast generation, with its mediation during this process seen as early as d2 of ESCs differentiation. While USP3-KD yielded decreased Flk1^+^ and Er71^+^ haemato-endothelial progenitors (∼ 50%) on d2 of ESCs differentiation, the reverse trend was seen on d5 (Fig. 6a, b). The same was further verified using USP3 efficient clones. While USP3-O/E led to an increase in the generation of both Flk1^+^ and Er71^+^ cells on d2 of ESCs differentiation, the converse was true on d5 when compared with that in control and USP3-mt clones (Fig. 6a, b). Immunostaining for Flk1 also confirmed the same (Fig. S5a). Together, our data suggested USP3 might be conferring a positive modulation during hemangioblast specification from ESCs *in vitro* and also expediting the same.

**Fig. 6.**
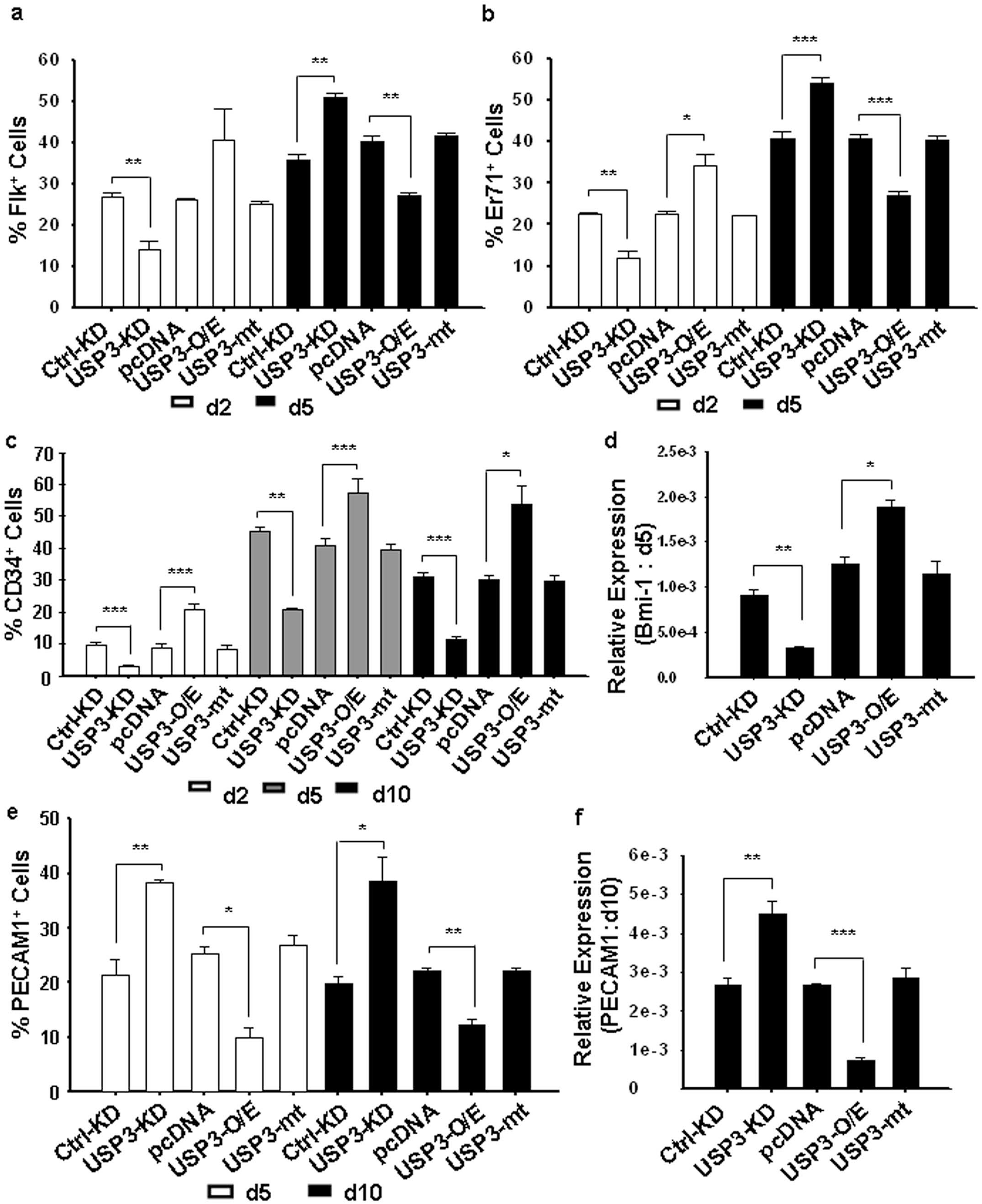
USP3 positively regulates Hemangioblasts specification and differentially influences haemato-endothelial differentiation. (a,b) Flow cytometry quantification of Flk1^+^ (a) and Er71^+^ (b) cells in USP3-KD and -O/E clones on d2 showed a positive modulatory effect of USP3 on hemangioblast specification, while the effect was reversed on d5 of ESCs differentiation; n = 3-4. (c) Flow cytometry quantification of CD34^+^ cells in USP3-KD and – O/E clones during ESCs differentiation on the stated time points showed a positive modulatory effect of USP3 on haematopoiesis; n = 3-7. (d) qPCR analysis for the expression of *Bmi-1* (d5 EBs), the HSCs marker, revealed a positive influence of USP3 on its expression; n = 3. (e) Flow cytometry quantification of Pecam-1^+^ cells representing the endothelial cells in d5 and d10 EBs in USP3 KD and -O/E clones revealed a negative modulatory effect of USP3 on endothelial specification; n = 3-4. (f) qPCR analysis of *Pecam-1* using d10 EBs in the stated clones also validated the negative influence of USP3 on endothelial differentiation from ESCs; n = 3. Data are means ± SEM. *, P ≤ 0.05; **, P ≤ 0.01; ***, P ≤ 0.001.

Further, we monitoread the downstream developmental events to assess the influence of USP3 during haematopoietic and endothelial differentiation from hemangioblasts. While USP3-KD decreased the CD34^+^, CD45^+^ and cKIT^+^ cells monitored at various time points (d2, d5 and d10) during mesodermal differentiation, and hence reflected the attenuation in haematopoietic differentiation, USP3-O/E promoted the same (Fig. 6c, Fig. S5d, e). The same was also further confirmed by immunostaining using various HSC-specific markers and also through expression analysis of Bmi-1 (∼ 2.7-fold reduction with USP3-KD, ∼ 1.5-fold increase with USP3-O/E and no difference in case of USP3-mt, when compared with respective controls) (Fig. 6d, Fig. S5c). Thus, a positive modulatory role of USP3 was evident during haematopoietic differentiation from hemangioblasts. On the contrary, USP3 negatively regulated endothelial differentiation, as discerned by monitoring the endothelial marker PECAM1 on d5 and d10 of mesodermal differentiation. While USP3-KD induced (∼ 2-fold) the PECAM1^+^ cells on both d5 and d10 of differentiation, the same was seen attenuated in USP3-O/E and with no alteration in USP3-mt, compared to the respective controls (Fig. 6e, f). Immunostaining for PECAM1 on d5 of ESCs differentiation also reinforced the same (Fig. S5a). Although, the population showing PECAM1 expression did not vary much from d5 to d10 under cardiomyogenic differentiation conditions; cells with USP3 deficient background generated endothelial cells with striking vessel-like morphology by d10 (Fig. S5b). Taken together, our study deciphered an interesting paradigm of differential cell fate modulation with USP3 at the cross-road to dictate haematopoietic and endothelial cell fates during early development. It displayed a positive modulatory effect directing induced hemangioblasts towards haematopoietic lineage, at the expense of endothelial ones.

### USP3, a histone DUB, mediates H2AUb/H2BUb deposition in the mesodermal genes

USP3 has been reported to function as a histone DUB and is involved in the DNA damage response pathway to protect genome integrity^26, 28^. Hence, to ascertain its chromatin association and functionality immunostaining was carried out using ESCs (data not shown) as well as the d5 EBs during differentiation. The detection of USP3 localization in d5 EBs with typical punctate pattern and colocalization with histone H2A within the nucleus proved it to be chromatin-associated, and hence, ascribed it as a chromatin-modulating enzyme (Fig. 7a). Next, we discerned its association with specific histone subunits and eventual plausible regulation of mesoderm differentiation as a histone DUB. Accordingly, control- and USP3-KD ESCs were differentiated and immunoprecipitation with USP3 was performed using d5 EBs during differentiation. Immunoblot for H2AUb and H2BUb revealed that USP3 interacted with both H2AUb and H2BUb (Fig. 7b). Similarly, reciprocal immunoblot for USP3 following pull down with H2AUb and H2BUb also confirmed the interaction between them during mesodermal differentiation (Fig. 7c). Intriguingly, the migration of the major interacting band, in case of both H2AUb and H2BUb immunoblots was seen similar to that of H2A and H2B (∼ 17kDa) respectively. We presume this might be due to the transiency of monoubiquitinated H2A and H2B, since their respective intensities were different from that of total H2A and H2B in each blot. The other distinct band was seen at ∼ 35kDa, probably reflecting the poly-ubiquitination status. Thus, both H2AUb and H2bUb were discerned to be the target of USP3 during ESC’s differentiation into the mesodermal lineage. The same was further strengthened by demonstrating increased levels of both H2AUb and H2BUb under USP3 deficient condition (Fig. 7d). On the contrary, USP3 efficient condition led to a decrease in the levels of H2AUb and H2BUb compared to that in USP3-mt and control (Fig.7d). Together these data reflected the importance of the catalytic activity of USP3 in regulating the histone deubiquitination dynamics. The same was further confirmed by quantifying the expression levels of H2AUb and H2BUb by western blot. As seen in Fig. 7e and f, the effect of USP3 on H2BUb (USP3-KD: ∼ 4-fold; USP3-O/E: ∼10-fold) was more striking compared to that of H2AUb (USP3-KD: ∼1.7-fold; USP3-O/E: 3-fold) when monitored on d5 of differentiation. Collectively, our findings emphasized on USP3 as a Wnt target functioning by regulating the turnover of histone ubiquitination, thereby determining the fate of mesoderm differentiation. To decipher the histone ubiquitin changes occurring in the promoter regions of *Bry, Mesp1, Pecam-1* and *Bmi-1* during mesoderm differentiation, we assessed the occupancy of H2AUb and H2BUb on the same. Chromatin immunoprecipitation (Chip)-qPCR analysis revealed occupancy of H2AUb and H2BUb on the promoter regions of the stated genes on d5 of ESCs’ differentiation (Fig. 7g). Notably, USP3-KD significantly increased the levels of repressive H2AUb in the *Bry* promoter (∼ 5-fold), whereas the active H2BUb mark was decreased (∼ 3-fold), corresponding with the reduction in the *Bry* transcription under USP3 deficient condition (Fig. 7g). On the contrary, USP3-KD significantly increased (∼ 2-fold) the occupancy of active H2BUb in the promoters of *Mesp1* corresponding to their increased expression under the USP3-KD background, while H2AUb remained unaffected. A similar trend in H2AUb and H2BUb occupancy (∼ 3-fold increase in H2BUb in USP3-KD) was also seen in the promoter region of PECAM-1 during endothelial differentiation in response to USP3. However, during haematopoietic differentiation, USP3-KD enhanced the levels of H2AUb and reduced the H2BUb mark by ∼3-fold each in the *Bmi-1* promoter (Fig. 7g) corroborating with the *Bmi-1* down-modulation under USP3-KD scenario. Overall, our study suggested a novel and context-dependent, both positive and negative modulatory influence of histone deubiquitination by USP3 during mesodermal fate choices and USP3 influencing the expression of mesodermal genes by differentially recruiting H2AUb and H2BUb (Fig. 8).

**Fig. 7.**
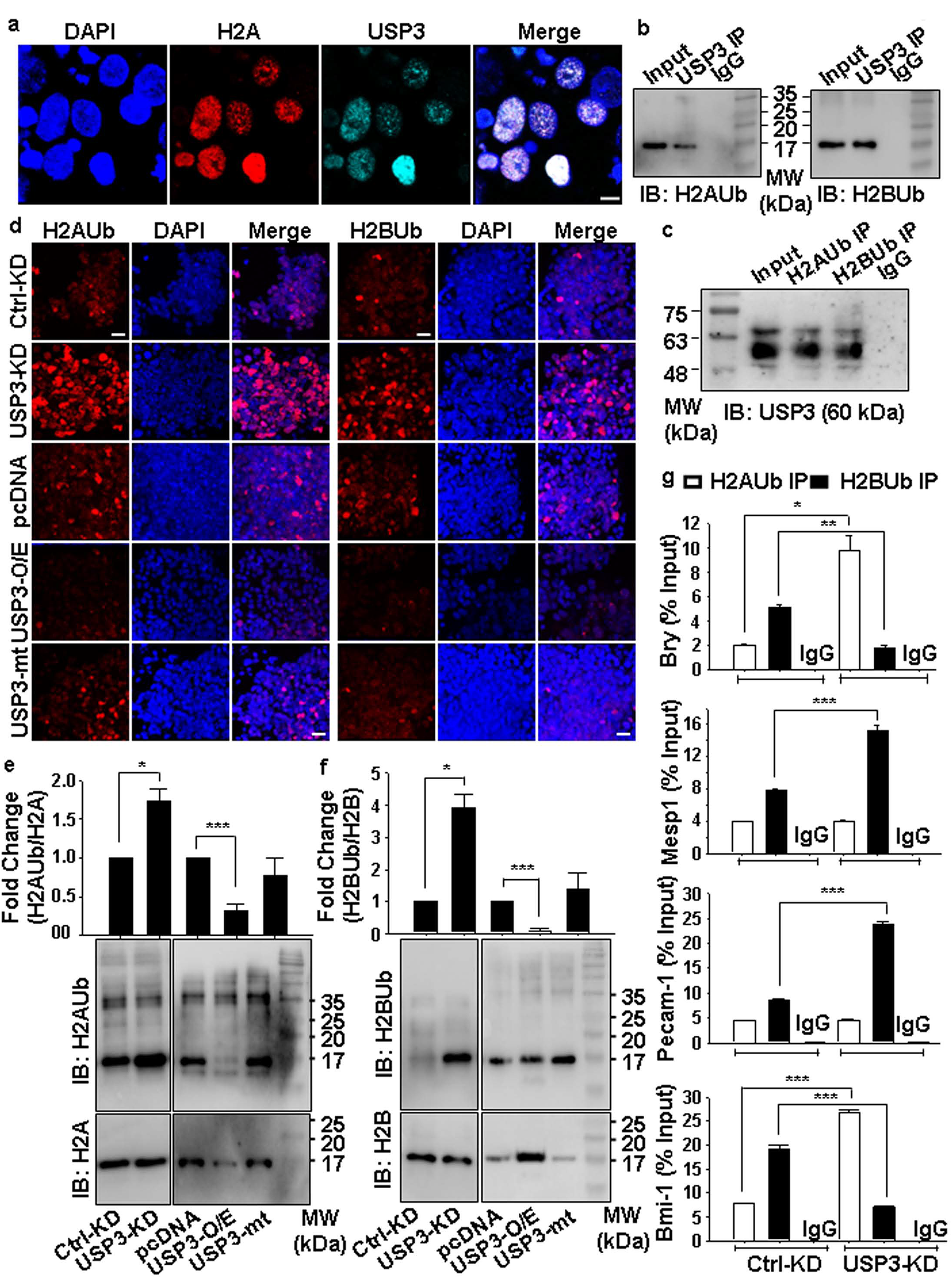
USP3, a histone DUB, mediates H2AUb/H2BUb deposition in the mesodermal genes. (a) Immunofluorescence analysis for USP3 in –LIF condition, carried out on d5 of ESCs differentiation, revealed nuclear localization of USP3 with punctate appearance. While the same was more evident with USP3-KD. (b, c) Immunoprecipitation demonstrates that USP3 binds to H2AUb and H2BUb. d5 EB Cell lysates were immunoprecipitated with anti-USP3 (b) or anti-H2AUb / anti-H2BUb (c), followed by immunoblotting with anti-H2AUb or anti-H2BUb for USP3 pull down or anti-USP3 for H2AUb/H2BUb IPs, respectively; n = 3-4. (d) Immunofluorescence analysis for H2AUb and H2BUb during d5 of ESCs differentiation with USP3-KD or USP3-O/E revealed histone deubiquitination activity of USP3 is required for H2A and H2B. (e,f) Western blot analysis for (e) H2AUb and (f) H2BUb with USP3-KD or O/E, revealed the deubiquitination activity of USP3 on H2A and H2B in d5 EBs during ESCs differentiation; n = 3. (g) Chromatin Immunoprecipitation followed by qPCR for H2AUb and H2BUb binding at Bry, Mesp1, Pecam-1 and Bmi-1 promoters displayed binding of H2AUb and H2BUb in the respective promoters. USP3-KD increased the occupancy of repressive H2AUb while reducing the binding of active H2BUb in the promoter region of mesoderm and hematopoietic-specific gene promoters. On the contrary, USP3-KD increased the occupancy of the H2BUb mark in the cardiac mesoderm as well as on endothelial-specific gene promoter. IgG was used as negative control and sonicated gDNA served as the Input control; n = 3. Data are means ± SEM; *, P ≤ 0.05; **, P ≤ 0.01; ***, P ≤ 0.001.

**Fig. 8.**
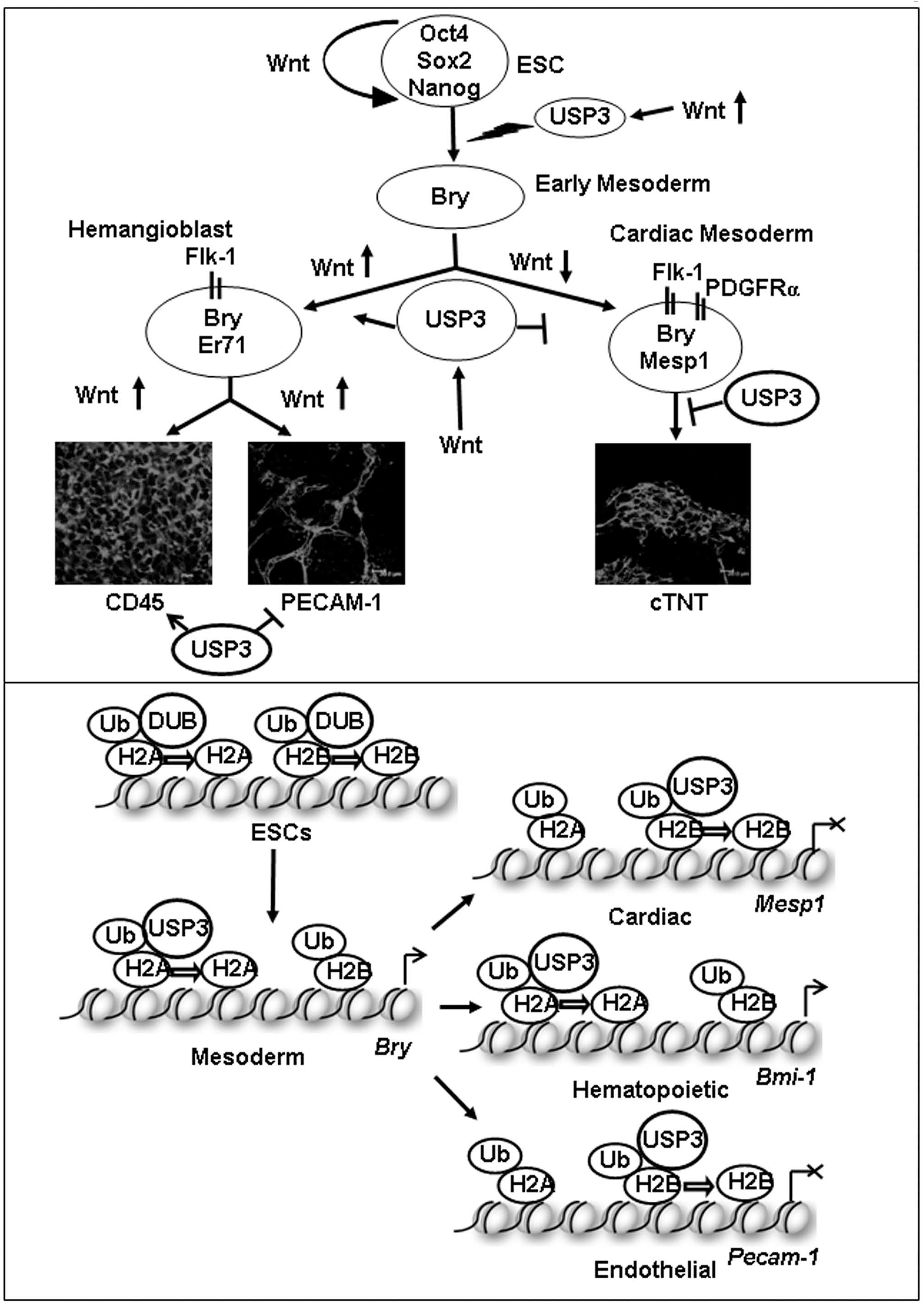
Schematic shows Wnt-USP3 link underlying early cell fate decision machinery during mesodermal specification and differentiation from ESCs (Upper panel); The both positive and negative modulatory influence of histone deubiquitination by USP3 during mesodermal fate choices via expression of mesodermal genes by differential recruitment of H2AUb and H2BUb (Lower panel).

## Discussion

The intricacies underlying early development do involve various genetic and epigenetic modulators during the transition of ESCs from undifferentiated to differentiated state^43–45^. Moreover, a delicate balance between ubiquitination and deubiquitination controls the cell fate decision machinery during embryonic development^15^. Over the years, several DUBs have been identified across species attesting various functional attributes during development. The data presented here establish the epigenetic modifier USP3, a histone DUB, as a crucial regulator of this transition. While USP3 was seen as dispensable during ESC’s maintenance, the USP3-dependent neuro-mesodermal lineage discrimination in differentiating ESCs did underscore the plausible role played by USP3 in the early lineage commitment process. Undoubtedly, this can be exploited further in the future to obtain lineage-specific differentiation from ESCs.

The involvement of several other DUBs has also been ascribed during neuro-mesoderm specification and differentiation. USP12 and USP46 are two such histone DUBs reported to have a modulatory role in mesoderm specification during *Xenopus* development^22^. A recent report has also suggested the expression status, but not the catalytic activity of USP12 influencing autophagy and neuroprotection^46^. While USP9X has been implicated during myogenesis using C2C12 skeletal myoblast cells^47^, Mysm1, USP10, USP16, and USP21 have been shown to be involved during haematopoiesis^48–52^, and USP7 and Mysm1 during adipogenesis and osteogenesis respectively^53, 54^. However, except for USP7^55^, a H3 DUB, none of these stated DUBs have been shown to be associated with Wnt signalling unlike that of USP3, a Wnt target gene as well as a modulator of Wnt signalling, as demonstrated by us in this report.

USP3 initially identified as a partial cDNA clone, contained highly conserved sequence regions common to all ubiquitin-specific proteases^15, 56, 57^. Further studies delineated USP3 as a chromatin-associated protein targeting ubiquitinated H2A/H2B/γH2AX and involved in regulating DNA damage response and maintenance of genome integrity^26, 28^. However, except for HSCs maintenance as revealed through cell cycle restriction and loss of chromosomal integrity in *USP3* null HSCs^27^, none of these reports suggested USP3 having any influence on cell fate decision. While this manuscript was in preparation, another recent article appeared suggesting USP3 involvement in the deubiquitination of and its interaction with OCT4 in human ESCs^58^. Notwithstanding, our findings unequivocally demonstrate the early developmental attributes of USP3.

Indeed, the expression profile of USP3 not only correlated well with that of *Bry* during *in vitro* ESCs differentiation to mesodermal lineage, but the presence of TCF/LEF binding motif in its promoter with the proven recruitment of β-catenin/TCF/LEF complex upon Wnt activation was also in congruence with that seen in *Bry* promoter. This is attributed to the association of USP3 and Bry with the gene regulatory network driving mesoderm differentiation. Moreover, the induction of USP3 in response to BIO at all the monitored stages during this differentiation suggested *USP3,* as a target gene of canonical Wnt signalling. USP4, USP7 and USP47 are the other USPs found to be important in preventing β-catenin degradation and stabilizing the same through their DUB activity and hence regulating Wnt signalling in various cancers^17, 18, 55^. However, in our study, we have demonstrated DUB having a dual function as a Wnt target and at the same time modulating the Wnt signalling and regulating Wnt responsive genes with profound early developmental relevance, a novel attribute for the Wnt-USP3 axis.

Wnt, a secreted glycoprotein, is a key developmental modulator having a profound influence on cell fate decision machinery^32^. Upon Wnt activation, TCF/LEF-1 binds to its consensus nucleotide sequences and together with β-catenin in the nucleus confers transactivation activity, and thus regulates the expression of target genes. During early embryogenesis, Wnt signalling is required for primitive streak development and mesoderm induction^32^. Also, Wnt proteins control multiple functions that include patterning of the anterior-posterior body plan, trunk/tail development, and specification of posterior mesodermal fate. Fate-mapping studies during mouse embryogenesis have shown that different mesodermal derivatives such as blood, vasculature, myogenic, renal tissues, etc. arise in an intricate and well-organized manner from distinct regions of the primitive streak^2^. Several reports including ours do suggest a temporo-spatial and context-dependent role of Wnt during mesodermal differentiation from ESCs^4, 6, 31, 59^. *Bry,* a ‘T box’ transcription factor, has been proven to be one such Wnt target gene that functions as a transcriptional activator genome-wide and forms a gene-regulatory feedback loop consisting of *Bry*, *Foxa2* and *Sox17* to orchestrate primitive streak formation in the early embryo^29, 60, 61^. Bry also facilitates the expression of canonical Wnt genes within the paraxial mesoderm precursors^60^. Using USP3 deficient and efficient conditions, our study demonstrated its indispensable role during mesoderm specification and further differentiation. While USP3-KD led to decreased expression of *Bry*, the same was seen up-regulated in USP3-O/E, but not in USP3-mt when monitored on d5 and d10 of ESCs differentiation. Our findings also suggested endogenous Wnt activation in USP3-O/E in d5 EBs was the major cause for enhanced mesoderm differentiation. Thus, a positive auto-regulatory loop might be functioning between USP3 and Wnt signalling in rendering mesoderm specification and the catalytic activity of USP3 might be important for the same.

Interestingly, USP3 also had a differential influence on the further differentiation of mesodermal cells to their derivatives. While USP3 was detected to be a positive modulator of hemangioblast specification (Flk^+^/Er71^+^), it negatively influenced cardiac progenitor specification (Mesp^+^/PDGFR^+^). Further, in line with our earlier report on Wnt influence during cardiomyogenic differentiation^4^, USP3 was also seen negatively regulating cardiomyogenesis. Indeed, the influence of USP3 corroborated well with that of Wnt influence during the specification of hemangioblasts and cardiac progenitors as well as further differentiation into cardiomyocytes (To be published elsewhere). Additionally, it displayed differential influence during haematopoiesis and endothelial differentiation from hemangioblasts. While USP3 was detected to be a positive modulator of haematopoietic differentiation (CD34^+^, cKIT^+^ and CD45^+^), it negatively regulated endothelial differentiation, as discerned by monitoring PECAM^+^ cells.

Endothelial-to-hematopoietic transition (EHT) is a dynamic process involving the shutting down of endothelial gene expression and switching on of hematopoietic gene transcription^62^. It is now widely accepted that RUNX1 is a key transcription factor involved in this process controlling the down-regulation and up-regulation of endothelial and hematopoietic transcriptional programs respectively^63^. USP3 was also seen to be inducing haematopoetic differentiation at the expense of endothelial differentiation suggesting its critical influence during the EHT process. Hence, it would be interesting to investigate further the association between USP3 and RUNX1 in materializing EHT.

Histone DUBs are the members that act upon the ubiquitinated histones, specifically H2AUb and H2BUb to remove their ubiquitination marks, rendering them either transcriptionally active (H2A) or repressive (H2B)^64^. In fact, monoubiquitinated H2A (H2AUb) and H2B (H2BUb) represent the dominant form of ubiquitinated histones^64^. Previous studies indicated that ubiquitination of both H2A and H2B regulates *Hox* gene expression and plays a critical role during embryonic development^65, 66^. Another study has suggested that the monoubiquitination of histone H2B on lysine 120 catalyzed by the E3 ligase RNF20 is increased during ESC’s differentiation and is required for the efficient execution of this process^23^. This increase is particularly important for the transcriptional induction of relatively long genes during ESC’s differentiation. Some of the recent studies have also shown that controlling the deubiquitination of H2BUb, which in turn controls gene repression, is critical in the transition from self-renewal to differentiation^23–25^. While USP16 regulates H2A deubiquitination and the gene expression in mouse ESCs during differentiation^25^, USP44 acts on H2BUb during the same^23^. Thus, there exist DUBs, highly specific for H2A and/or H2B and contributing immensely during early development. Interestingly, USP3 has been reported to regulate transcription by deubiquitinating both histone H2A and H2^26, 28^. Our study has also demonstrated USP3 functioning as a histone DUB with similar specificity for both H2A and H2B during mesoderm differentiation. Moreover, USP3 not only regulated *Bry* expression during d2 and d5 of ESCs differentiation but also modulated mesodermal cell fate by orchestrating histone deubiquitination dynamics in the *Bry* promoter. While USP3-KD was seen up-regulating both H2AUb and H2BUb, its over-expression reversed the same during ESCs differentiation into Bry^+^ mesodermal cells. A similar finding was also reported in *Xenopus*^22^, where USP12 specifically regulated the expression of mesoderm marker, *Xbra*, during the gastrula stages of developing embryo with H2AUb enrichment in the promoter regions of *Xbra*. However, H2BUb was not detected in the *Xbra* promoter and USP12 knockdown led to an increase in the H2B ubiquitination. In contrast, we observed the presence of both H2AUb and H2BUb in the *Bry* promoter on d5 of ESCs differentiation. Moreover, USP3-KD association with increased occupancy of H2AUb and decreased H2BUb indeed correlated well with diminished *Bry* expression on d5 in USP3-KD. USP3 also displayed differential enrichment of H2AUb or H2BUb in the promoters of mesodermal derivative genes. In contrast to the reduction in the transcription of the HSC marker, *Bmi-1,* with USP3-KD that was associated with enhanced occupancy of H2AUb, the upregulation of cardiac and endothelial marker genes was correlated with the enrichment of H2BUb. Mysm1 is the other DUB, which was shown to have similar specificity to H2A and H2B and plays an important role during haematopoietic development^67^.

Indeed, our study has added USP3 and histone deubiquitination to the long list of factors regulating mesoderm differentiation. Our study has not only provided further evidence for H2AUb and H2BUb in gene regulation but also elucidated a more comprehensive understanding of the functions of histone ubiquitination in chromatin and cellular regulation. It seems likely that, DUBs like USP3 might distinguish the respective target gene loci via plausible differential interaction with the sequence-specific transcription factors that recruit them across the genome. Exploring further the underlying basis on how USP3 functions selectively as either a co-activator or co-repressor of gene transcription during early development would facilitate unleashing the associated key modulators in dictating the cell fate decision machinery in a temporo-spatial manner, in parallel to revealing other plausible functions that USP3 may have during the same.

## Materials and Methods

### ESCs’ maintenance and differentiation

All experiments were carried out using murine ESC line, D3 (ATCC, Manassas, VA), and the derived stable clones for Mesp1-Venus (the plasmid was a kind gift from Dr. Blanpain)^41^. ESCs were maintained in culture using Dulbecco’s modified Eagle medium (DMEM) with leukemia inhibitory factor (LIF) (1,000 U/ml; Merck-Millipore), L-glutamine, penicillin-streptomycin, nonessential amino acids (all from Invitrogen, Grand Island, NY), and β-mercaptoethanol (Sigma-Aldrich, St. Louis, MO) and were passaged every 48 h. Differentiation of ESCs into cardiomyocytes was induced by generating embryoid bodies (EBs) in hanging drop (500 cells/20 μl drop) for 2 days (d0-2) followed by suspension culture for 3 days (d2-5) using non-adhesive dishes as reported earlier^4^. The medium used for differentiation was the same as the maintenance medium *sans* LIF. EBs were plated on gelatine-coated tissue culture dishes (24-well dish; 1EB/ well) on d5 and were monitored at various stages of development. Cardiomyogenic differentiation efficacy was discerned by counting (i) the number of beating EBs, and (ii) beating clusters/EB on day 5 following their plating, i.e., d10 of cardiac differentiation. To study the effect of Wnt signalling, BIO (6-Bromoindirubine-3’-oxime; Calbiochem, La Jolla, CA, USA) (1µM), and DKK1 (R&D, Minneapolis, MN, USA) (50ng/ml) were added, as indicated^4^. DMSO served as the vehicle control for BIO.

### Transfection and generation of loss-of-function and gain-of-function stable ESC clones

Two sets of USP3 shRNAs were designed against its target sequence and cloned independently into pSilencer2.1-U6 hygro vector (Ambion). Scrambled shRNA with limited homology to any known sequences was cloned into the same vector and could serve as the negative control. Stable USP3 knockdown (USP3-KD) and scrambled (Ctrl-KD) clones were generated by nucleofection (Amaxa) using Mesp-Venus ESCs, following the manufacturer’s instructions. Similarly, stable ESC clones were generated overexpressing USP3 (USP3-O/E) and its catalytic mutant (Myc-USP3C168S, designated as USP3-mt) (Both vectors, kind gift from Dr. Citterio, Netherlands Cancer Institute)^26^ under the D3 wild type ESCs backdrop and with pcDNA, the empty vector, serving as the control. USP3 harboured two conserved protein domains: a catalytic domain of the Ub-specific protease (USP) class and a zinc finger (ZnF-UBP) Ub-binding domain. USP3-mt vector had a substitution of serine^168^ with a cysteine residue, thus making it catalytically inactive^26^.

### RNA isolation and transcript analysis

Total RNA was isolated using TriReagent (Sigma) and was processed further for DNAse treatment followed by reverse transcription-PCR. Quantitative PCR (qPCR) was performed using SYBR green supermix with a CFX thermocycler (Bio-Rad, New South Wales, Australia) following the conventional protocol. Detail primer sequences are given in Table 1 as a supplementary file. Relative expression levels of each gene were analyzed after normalizing with β-actin and the fold difference was calculated by comparing that with the control.

### Immunocytochemistry

The cellular phenotypes were studied by immunocytochemistry, following the standard protocol. In brief, cells were fixed with 4% paraformaldehyde (PFA) followed by permeabilization, as necessary, using 0.25% Triton X-100 (Sigma) and blocking with 5% fetal bovine serum (FBS) in phosphate-buffered saline (PBS). The cells were then incubated with respective primary antibodies, as desired, followed by incubation with Cy3-conjugated (Chemicon) secondary antibodies for fluorescent labelling. 4,6-Diamidino-2-phenylindole (DAPI) (Sigma) was used as a nuclear counterstain. In each case, the negative control was performed with the replacement of respective primary antibodies with FBS. The slides were observed under the laser-scanning confocal microscopes (TE2000E; Nikon, Leica SP5 II, or Zeiss LSM510) to detect the labelled cells and the cellular localization of the stated candidates.

### Histone staining

ESCs or the EBs were resuspended in hypotonic buffer (10 mM Tris-HCl, pH 7.5, 10 mM NaCl, 5 mM MgCl2) and incubated at 37°C for 15 min. 1/10 volume of 3% Tween 20 were added to the cell suspension and incubated for 5 min. Subsequently, the cells were washed and incubated with KCM-T solution (10 mM Tris, pH 8.0, 120 mM KCl, 20 mM NaCl, 0.5 mM EDTA, 0.1% Tween 20) containing 0.1% Triton X-100 at RT for 10 min. After 3x washing with KCM-T buffer (without Triton X-100), the cells were fixed using 4% PFA at RT for 10 min followed by further washing with KCM-T buffer (3x). The samples were blocked with KCM-T buffer containing 2.5% BSA and 10% serum at RT for 1 h. Further, the samples were incubated with primary antibody at RT for 4 h or at 4 °C O/N. After thorough washing with KCM-T buffer, the cells were incubated with secondary antibodies and DAPI for an additional 1 h. Finally, the cells were thoroughly washed with KCM-T buffer (3x) followed by PBS washing (2x) and mounted on a glass slide using Vectashield mounting medium. The slides were observed under a laser-scanning confocal microscope, as described earlier.

### Flow cytometry quantification

For flow cytometry quantification, a single-cell suspension was prepared and the cells were incubated with primary antibodies for 45 min, followed by washing and incubation with secondary antibodies for 30 min, as described^3^. Both the primary and secondary antibodies were suspended in a staining solution containing 2% FBS and 0.05% saponin in PBS, and the incubations were carried out at 4 °C. Immunostaining of the single-cell population was performed using primary antibodies followed by phycoerythrin (PE)-conjugated secondary antibodies (BD Biosciences, San Jose, CA) for fluorescent labeling. Quantification was carried out using a FACS Canto instrument (Becton Dickinson, Singapore). About 10,000 viable cells were analyzed per sample, following acquisition settings with appropriate controls. The emitted fluorescence of Mesp-venus and PE were measured on a log scale at 530 nm and 585 nm, respectively. The analysis was performed using the FACS Diva software program (Becton Dickinson).

### Immunoprecipitation

For immunoprecipitation experiments, cells were harvested using Lysis buffer (50mM Tris-HCl; pH. 7.5, 5mM EDTA, 250mM NaCl, 50mM NaF, 1mM sodium orthovanadate, 0.5% Triton X-100, 1mM Phenylmethylsulfonyl fluoride and 1X protease inhibitor cocktail). The protein concentration of the whole cell lysate was estimated using Biorad Protein Assay Dye reagent using BSA (0.1mg/ml) as standard. Cell lysates (500 μg) were incubated with 2 μg of the indicated antibodies at 4 °C overnight. Dynabeads Protein A/G (Dynal, Invitrogen) were washed in 3% BSA/1x PBS and incubated with the cell lysate for 2 h. The immunoprecipitates were washed and proteins were eluted by boiling in the loading buffer. The resulting protein samples were resolved using SDS PAGE followed by transfer onto the PVDF membrane. Blocking was carried out with either 5% BSA or 5% non-fat dried milk followed by incubation with respective primary antibodies and then with secondary antibodies conjugated with HRP (Biorad). Blots were developed using Western Bright Peroxide chemiluminescent detection reagent (Advansta) and documented. Densitometry-based analysis was performed using Quantity One software.

### Chromatin Immunoprecipitation

Chromatin preparation and ChIP were essentially performed, as described^68^. Briefly, d5 EBs were trypsinized and the single cells thus obtained were cross-linked with 1% formaldehyde in PBS for 20 min at room temperature, followed by quenching by the addition of 125 mM glycine for 5 min on ice. After 2x PBS wash, the cross-linked cells were washed sequentially using the buffers, as state^68^. Subsequently, cells were disrupted using a sonicator (Bioruptor, Diagenode) to produce an average DNA fragment size of 250 bp. The cell lysates were incubated with desired antibodies (β-catenin, H2AUb, H2BUb) at 4 °C overnight. Dynabeads Protein A/G (Dynal, Invitrogen) were equilibrated with 3% BSA/1x PBS and incubated with the cell lysate for 2 hours at 4 °C. Antibody ChIPs were further washed using buffers containing increasing concentrations of NaCl followed by no NaCl and finally eluted by adding 400 µl elution buffer containing 0.1 M NaHCO3 and 1% SDS^68^. Reverse crosslinking of the ChIP samples was carried out at 65°C overnight using NaCl. For DNA purification, the ChIP samples were treated with RNAse followed by proteinase K by incubating for 2 hrs each at 37°C. DNA was extracted using phenol-chloroform followed by ammonium acetate/ ethanol precipitation. Subsequently, samples were analyzed by qPCR for the promoter regions of USP3 (-180 to +60), Brachyury (-260 to -110), Mesp1 (-200 to +5), Pecam-1 (-200 to +45), and Bmi-1 (-720 to -440).

### Teratoma induction and Immunohistochemical analysis

All the animal experiments were conducted as per the Institutional Animal Ethical Committee guidelines and approval. Around 0.2 x10^6^ ESCs (in 100µl PBS) were injected subcutaneously into the dorsal flanks of 4–6-week-old male NOD/SCID mice. Induction of teratoma was seen by 4-5 wk of cell injection. The mice were sacrificed by cervical dislocation, tumours were excised, and fixed in 4% PFA overnight followed by overnight sequential incubation with 15% and 30% sucrose gradient each. After embedding those in Cryomatrix, frozen sections (20µm) of tumours were taken using cryotome. Tissue sections were subjected to antigen retrieval using 10mM sodium citrate buffer followed by heating for 2 min and then cooling subsequently. Permeabilisation and blocking were carried out using 5% FBS and 0.8% BSA prepared in PBS containing 0.3% Triton X-100 for 4hr at room temperature. Sections were then stained for three germ layer-specific markers; ectoderm: nestin (Chemicon), endoderm: AFP (Santacruz), and mesoderm: brachyury (Santacruz) respectively by incubating with the stated primary antibodies at 4 ^ο^C overnight. Excess antibody was washed off using PBS, followed by incubation with secondary antibodies conjugated with Cy3 for an hour at room temperature in the dark for fluorescent labelling along with DAPI as the nuclear counterstain. Slides were then imaged using the confocal laser-scanning microscope to detect fluorescence.

### Statistical analysis

All quantitative data were presented as mean ± S.E.M (standard error of the mean) and statistical significance was calculated using either paired or unpaired Student’s “t*”* test (Sigmaplot, San Jose, CA). “P” values were calculated in comparison with the control and were represented as follows: *, P ≤ 0.05; **, P *≤* 0.01; *\*\*\**, P *≤* 0.001. All of the statistical analyses performed in this study and their respective parameters are shown in the figure legends. Source data are available online. The number (*n*) of biological repeats is indicated in each figure legend. All representative data are shown from independently repeated experiments or independent animals with similar results.

## Acknowledgement

The work was supported by funding support from DBT (BT/PR16655/NER/95/132/2015) and Indo-Australia Biotechnology Fund (BT/Indo-Aus/06/02/2011) to NL. VH wishes to acknowledge the fellowship from the Council of Scientific and Industrial Research, India received towards his graduate studies.

## Author contribution

Conceptualization and Resource generation: NL, Execution: VH, Analysis and Interpretation: VH and NL, Manuscript writing and finalization: NL and VH.

## Supplementary data

**Fig. S1: USP3 doesn’t influence ESC characteristics during maintenance.** (a-c) qPCR analysis showing no significant difference in the expression of (a) *Sox2* (b) *cMYC* (c) *Klf4* in USP3-KD and -O/E clones in comparison with their respective controls. Data were presented as fold difference with respect to control values set at 1; n = 3. (d) Immunofluorescence analysis for Oct4, Nanog and Sox2 in USP3-KD and -O/E clones showing similar expression of these pluripotency-associated markers when compared with their respective controls. Scale bar: 20μm.

**Fig. S2: Influence of USP3 during spontaneous differentiation of ESCs.** Immunofluorescence analysis for Oct4 and Nanog in ESCs grown under +LIF and –LIF conditions for 5 days revealed their comparable expression in USP3-KD and -O/E clones similar to that of respective controls, Scale bar: 20μm.

**Fig. S3: Teratoma induction and assessment of USP3 pluripotency *in vivo*.** Ctrl-KD, USP3-KD, pcDNA and USP3-O/E ESCs were able to generate teratomas in immuno-compromised mice and contained cells of all three germ layers. Immunostaining with Nestin, Brachyury and AFP was performed to observe the cells of mesoderm, endoderm and ectoderm respectively. While USP3 displayed mesoderm-promoting and ectoderm-suppressing influence, endoderm remained unaffected. Scale bar: 20μm.

**Fig. S4. The role of USP3 during cardiomyogenic specification and differentiation from ESCs *in vitro*.** (a) Immunofluorescence analysis for Mesp1 expression representing the cardiomyogenic progenitors on d5 of ESCs differentiation displayed the negative influence of USP3 on their generation; Scale bar: 20μm. (b, c) qPCR analysis for the expression of cardiac markers (b) Mef2c and (c) Mlc2a in d10 EBs revealed an increase in their expression with USP3-KD, while the reverse was true in the case of USP-O/E and no alteration in USP3-mt when compared with their respective controls; n = 3. Data are means ± SEM; *, P ≤ 0.05; **, P ≤ 0.01; ***, P ≤ 0.001.

**Fig. S5.** The role of USP3 during hemangioblast specification and further differentiation to hematopoietic and endothelial derivatives. (a, b) Immunofluorescence analysis for (a) Flk-1 and Pecam-1 expression on d5 of ESCs differentiation revealed a negative influence of USP3 on the generation of endothelial progenitors. (b) The same was also reflected by observing endothelial cells with striking vessel-like morphology on d10 of differentiation in USP3-KD cells compared to control (Ctrl-KD). (c) On the contrary, USP3 rendered a positive influence on haematopoiesis as ascertained by immunofluorescence detection of CD34 and CD45 expression on d5 of ESCs differentiation. Scale bar: 20μm. (d, e) The positive influence of USP3 was further validated by quantifying CD45^+^ (d) and cKIT^+^ (e) cells at various time points of ESCs differentiation. Data are means ± SEM; n = 3-7. *, P ≤ 0.05; **, P ≤ 0.01; ***, P ≤ 0.001.

