## Supplementary figures and images for "Orchestration of differential mesodermal fate choice from ESCs by Wnt-USP3 link and H2A/H2B contextual deubiquitination"

### Suppl. Fig. S1-S5

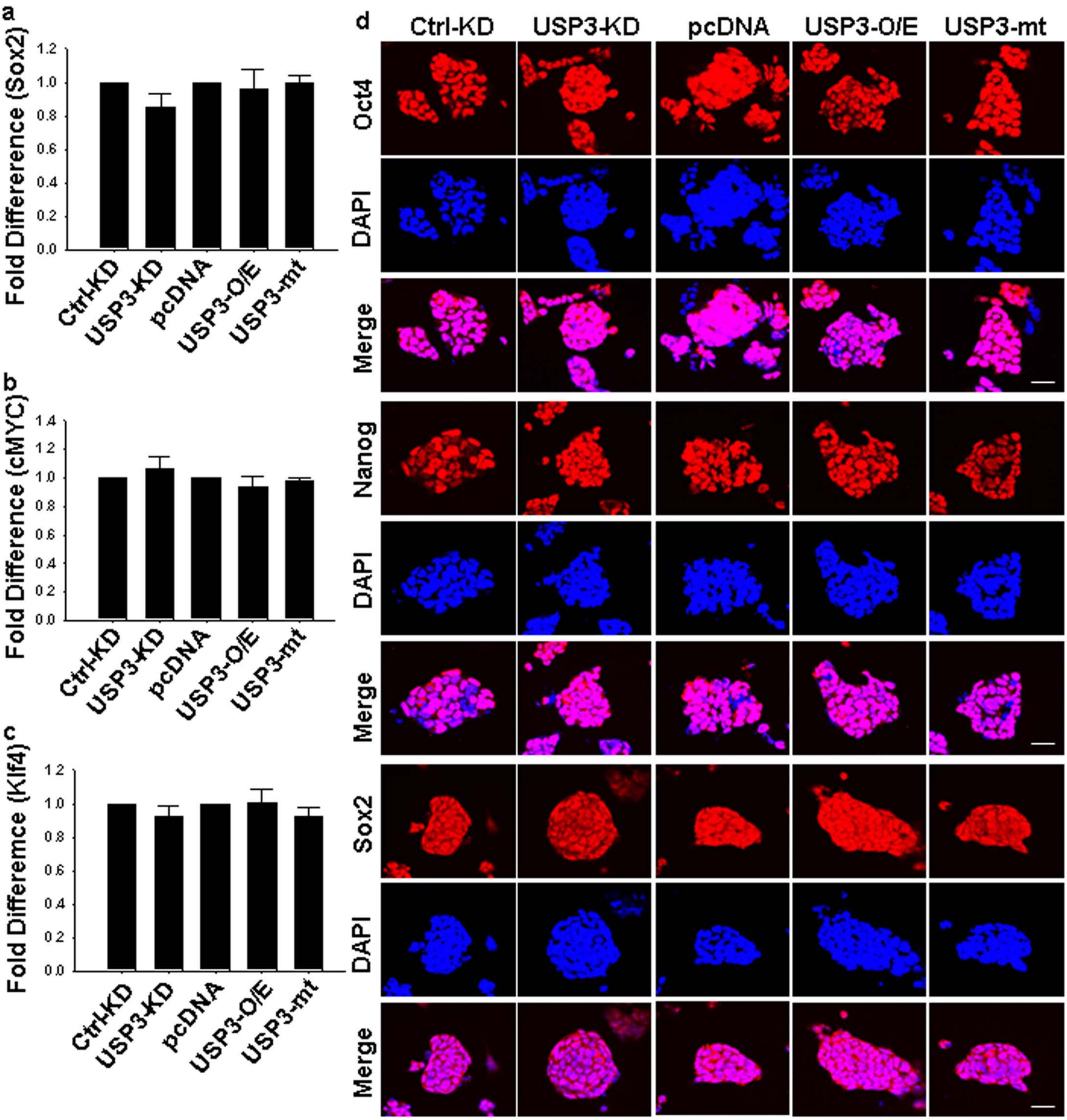

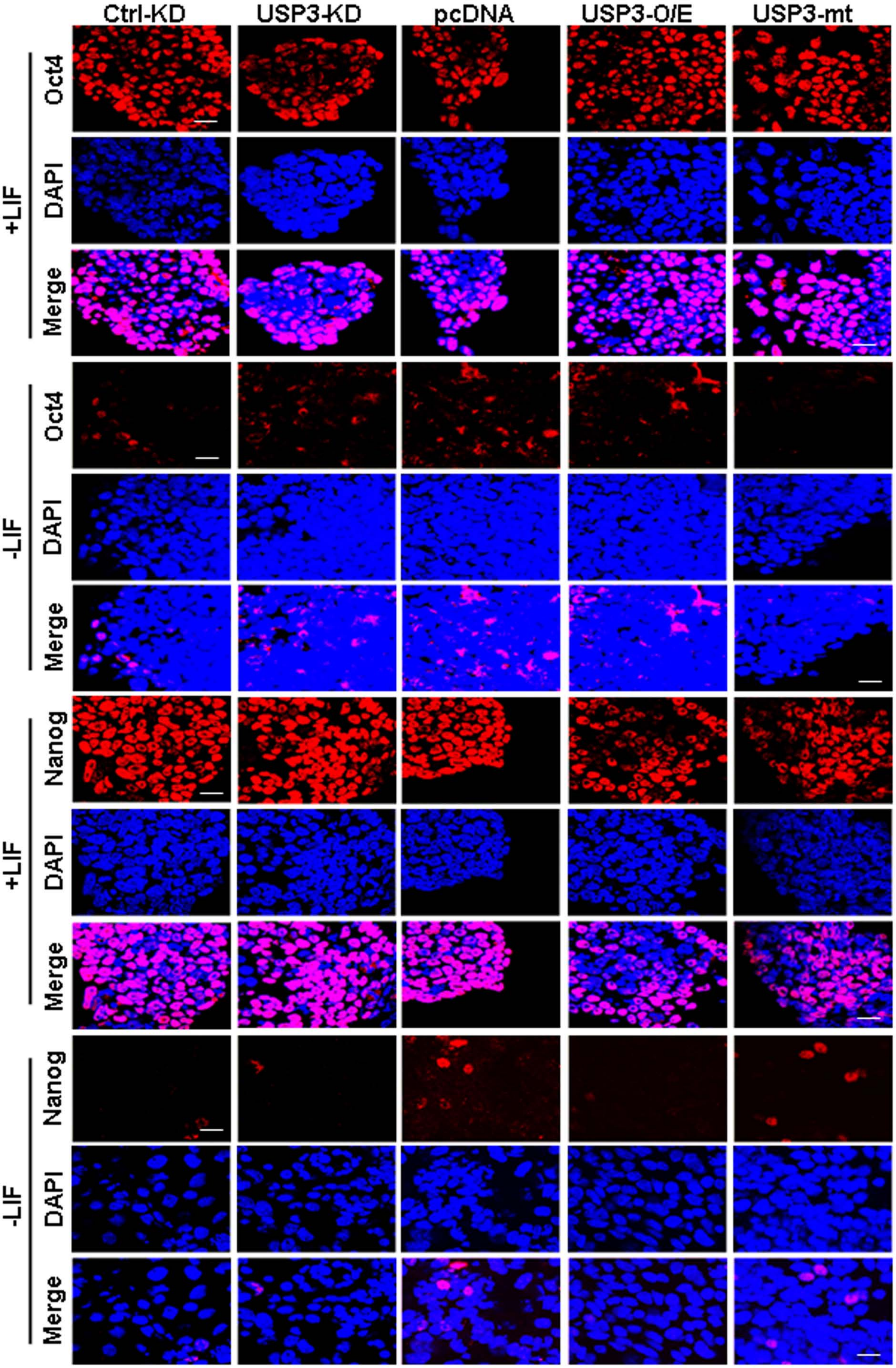

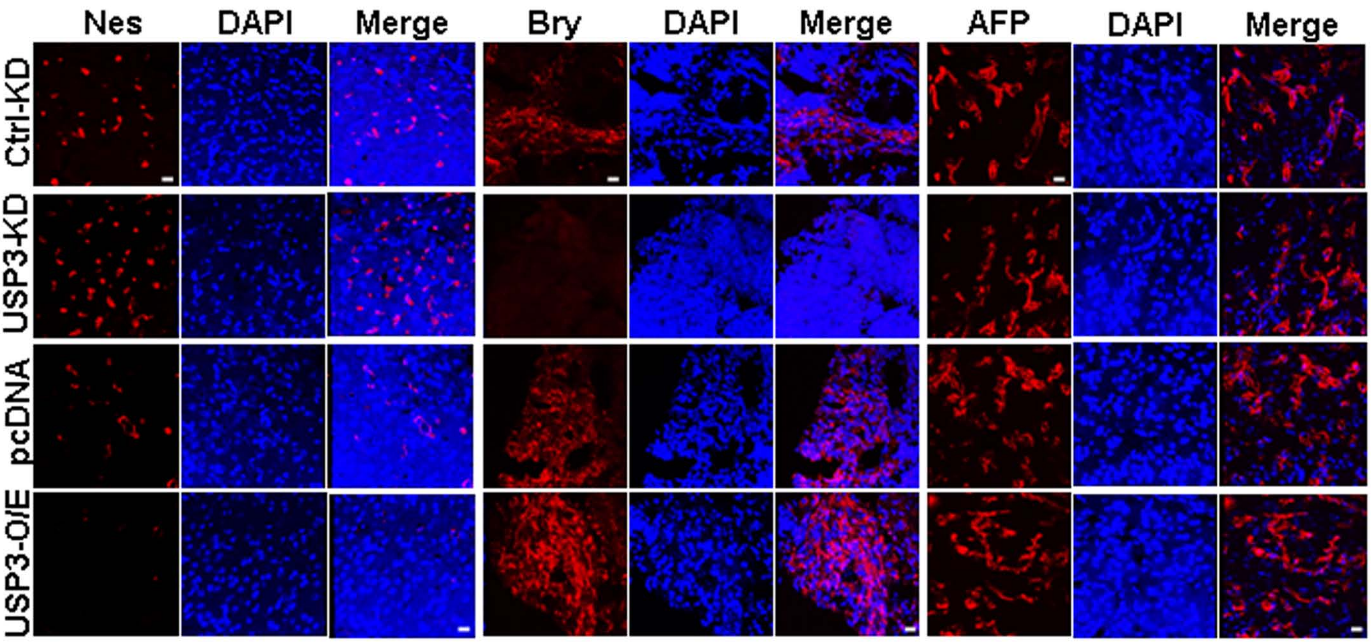

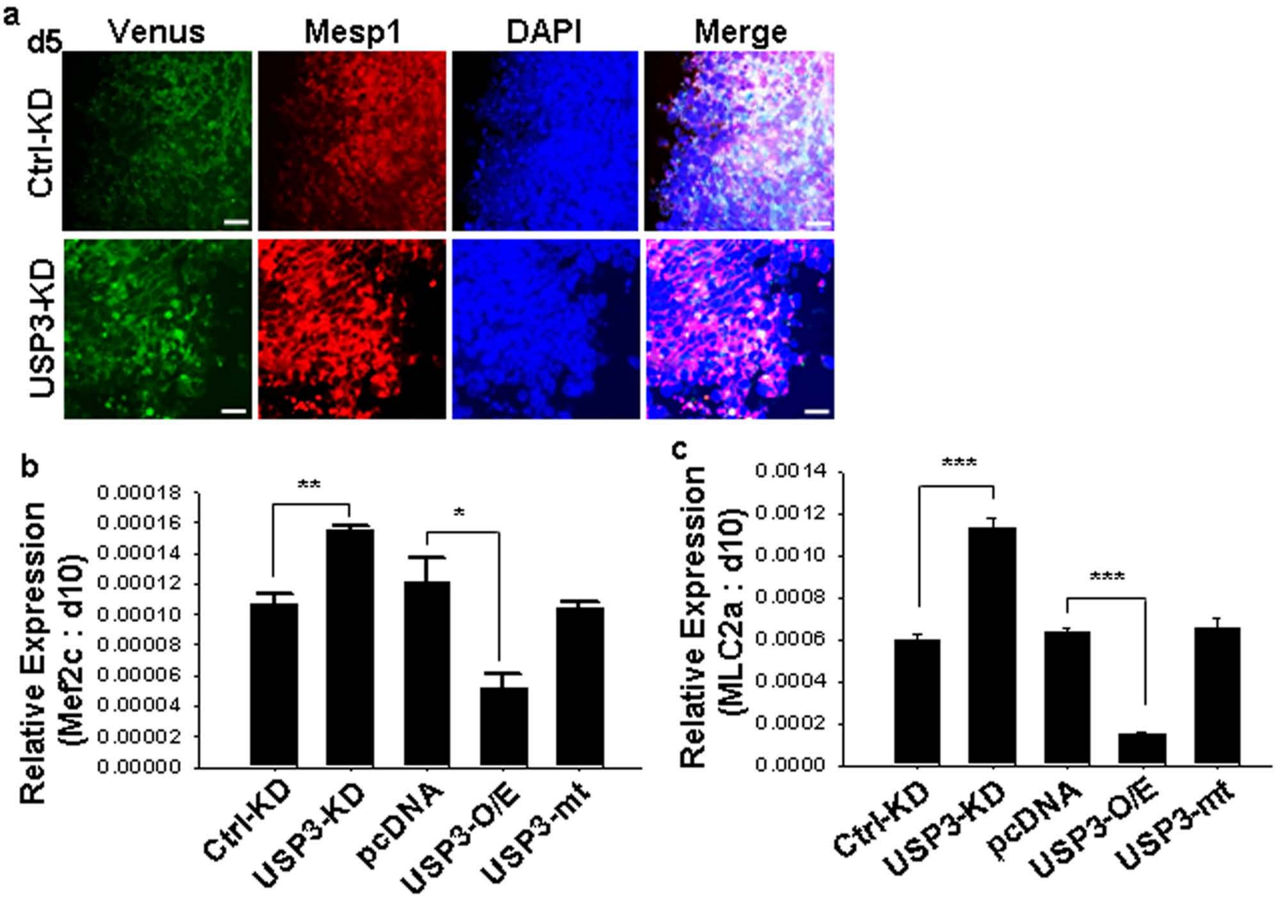

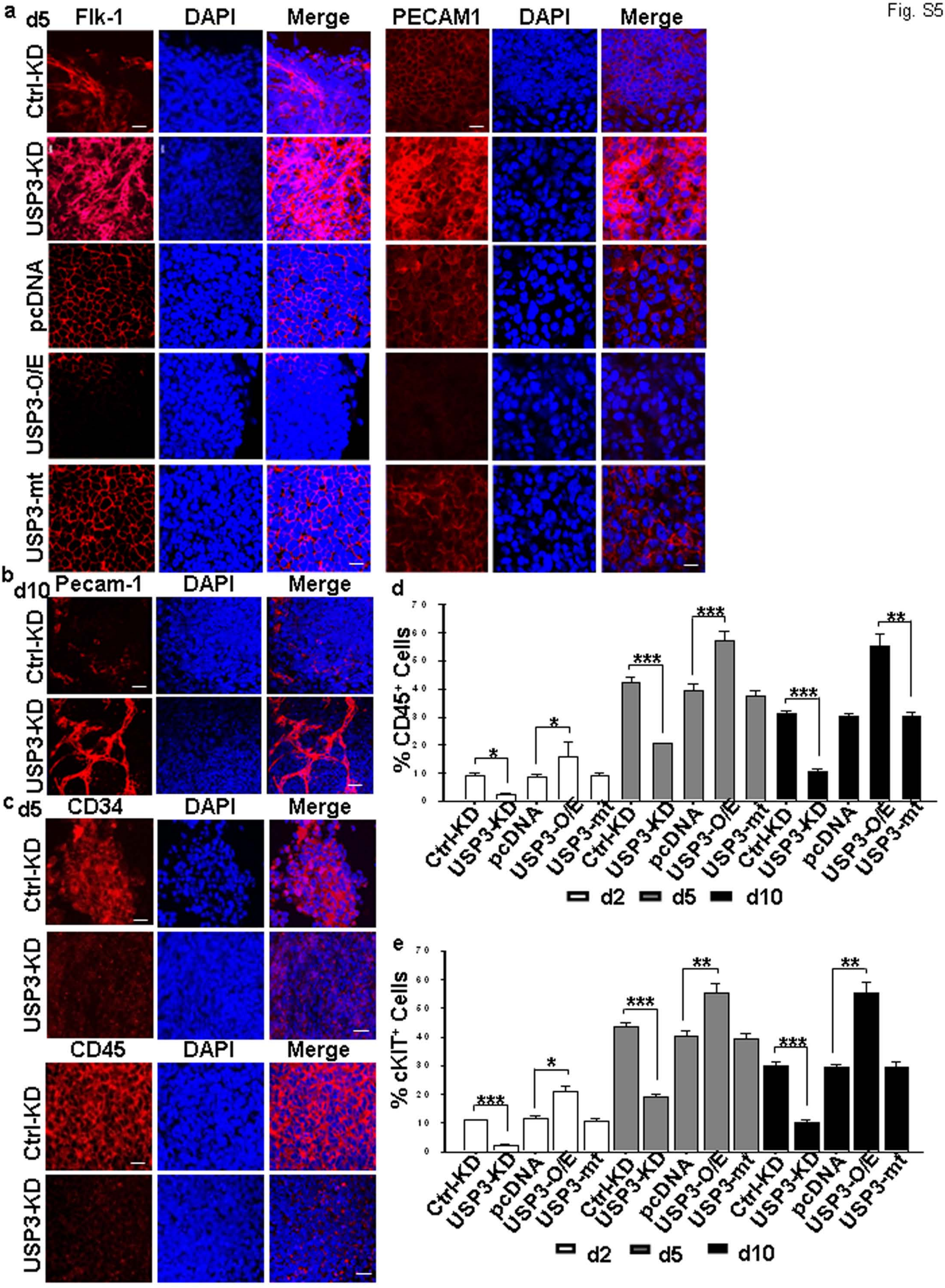
